# The intracellular auxin homeostasis regulators PIN5 and PIN8 have a divergent membrane topology in *Arabidopsis thaliana* root cells

**DOI:** 10.1101/2022.12.27.522031

**Authors:** Yewubnesh Wendimu Seifu, Nikola Rýdza, Marta Zwiewka, Vendula Pukyšová, Tomasz Nodzyński

## Abstract

PIN proteins establish the auxin concentration gradient, which coordinates plant growth. PIN1-4 and 7 localized at the plasma membrane (PM) and facilitate polar auxin transport while the endoplasmic reticulum (ER) localized PIN5 and PIN8 maintain the intracellular auxin homeostasis. Although an antagonistic activity of PIN5 and PIN8 proteins in regulating the intracellular auxin homeostasis and other developmental events have been reported, how the two proteins which localize at the same intracellular compartment antagonize each other remains unclear. Combining immunolocalization, pH-dependent fluorescent quenching, and topology prediction programs, we mapped the membrane topology of PIN5 and PIN8 in *Arabidopsis thaliana* root cells. Our results indicate that, except for the similarities in the orientation of the N-terminus, PIN5 and PIN8 have an opposite orientation of the central hydrophilic loop and the C-terminus, as well as an unequal number of transmembrane domains (TMDs). PIN8 has ten TMDs with groups of five alpha-helices separated by the central hydrophilic loop (HL) residing in the ER lumen, and its N- and C-terminals are positioned in the cytoplasm. However, topology of PIN5 comprises nine TMDs. Its N-terminal end and the central HL face the cytoplasm while its C-terminus resides in the ER lumen. Overall, the divergent membrane topology of PIN5 and PIN8 provides a possible explanation for the mutually opposing activity of these intracellular auxin homeostasis regulators.

## Introduction

Auxin regulates plant growth and development through its concentration gradient, which is established by the activity of various auxin transporter proteins including auxin influx carrier AUX/LAX family and the auxin efflux PIN-FORMED (PIN) proteins (van Berkel *et al*., 2012). The PIN-FORMED (PIN) proteins are a family of integral membrane proteins found in almost all land plants (Viaene *et al*., 2014; Mutte *et al*., 2018). *Arabidopsis thaliana* genome encodes eight PINs which are classified into the plasma membrane (PM) localized PINs (PIN1, 2, 3, 4 and 7), the endoplasmic reticulum (ER) localized PINs (PIN5, and 8), and the dual PM- and ER-localized PIN6 (Mravec *et al*., 2009; Ding *et al*., 2012; Nodzyński *et al*., 2016; Simon *et al*., 2016; Zwiewka *et al*., 2019; Sisi & Růžička, 2020).

The topology of PIN proteins is composed of trans-membrane domains (TMDs) separated by a central hydrophilic loop. The structure of the TMDs is highly conserved while the hydrophilic loop is varied in size and amino acid composition (Zwiewka *et al*., 2019; Sisi & Růžička, 2020). The PM PINs have a longer central hydrophilic loop, the ER PIN5 and PIN8 have a shorter loop (Mravec *et al*., 2009; Ganguly *et al*., 2010), and PIN6 has an intermediate sized loop (Simon et al., 2016). The central hydrophilic loop (HL) of the PM PINs localizes in the cytoplasm (Nodzyński *et al*., 2016), and contains various motives, such as phosphorylation sites which modulate the polar PM distribution of the PINs (Zourelidou *et al*., 2014; Xi *et al*., 2016; Zwiewka *et al*., 2019). The subcellular polarity of the PM PINs is essential to mediate the directional cell-to-cell auxin transport. Conversely, the ER-localized PIN5 and PIN8 do not have phosphorylation sites, and their role is to mediate the intracellular auxin homeostasis (Mravec *et al*., 2009; Ding *et al*., 2012; Sisi & Růžička, 2020).

Studies have shown that PIN5 and PIN8 proteins have antagonistic activities in regulating the intracellular auxin homeostasis and various developmental aspects (Mravec *et al*., 2009; Ganguly *et al*., 2010; Ding *et al*., 2012). Auxin transport assay utilizing protoplasts isolated from a PIN5 overexpressing line revealed a decreased indole-3-acetic acid (IAA) export activity, whereas *pin5* knock-out mutants exhibited a higher level. Therefore, it was proposed that PIN5 conveys auxin from the cytoplasm to the ER lumen (Mravec *et al*., 2009). In contrast, analogue study using PIN8 overexpressing and *pin8* knock-out lines showed an increased and a decreased auxin export activity respectively, indicating that, antagonistically to PIN5, PIN8 may transport auxin from the ER lumen into the cytoplasm (Ding *et al*., 2012). In addition, the PIN5 overexpressing line showed a higher accumulation of IAA conjugates and a lower level of free IAA (Mravec *et al*., 2009). However, the PIN8 overexpressing plants exhibit a higher level of free IAA and a decreased accumulation of its conjugates (Ding *et al*., 2012). Furthermore, the opposing activities of these proteins were demonstrated in the analysis of root hair growth (Ganguly *et al*., 2010), lateral root development (Lee *et al*., 2020), hypocotyl growth (Ding *et al*., 2012), and leaf venation pattern (Sawchuk *et al*., 2013).

Although the opposite developmental roles of the ER-localized PIN5 and PIN8 and their countering influence on the intracellular auxin homeostasis have been reported, the reason why these proteins oppose each other remains unclear (Ding *et al*., 2012), and the membrane topology of both proteins is poorly understood. Elucidating the membrane topology of these proteins, their orientation in reference to the membrane within which they reside, is essential to understand whether the contradicting activities of these ER PINs might be related to their structural differences. Therefore, in this study, we investigated the membrane topology of PIN5 and PIN8 in *Arabidopsis thaliana* root cells in terms of the sub-cellular orientation of their central hydrophilic loop, and the N- and C-terminal ends. Our data show that, except for the similarities in the orientation of the N-terminus, PIN5 and PIN8 have opposite subcellular orientations of the central hydrophilic loop and the C-terminal end. These findings indicate that the mutually opposing activity of PIN5 and PIN8 could be attributed to their topological differences.

## Materials and Methods

### Plant Material and Growth Conditions

The previously published *Arabidopsis thaliana* transgenic lines: PIN2::PIN5-GFP, and C1 (PIN5 chimeric protein which contains GFP fused PIN2-HL in the PIN5-HL) (Ganguly *et al*., 2014), 35S::PIN8-GFP (Ding *et al*., 2012), 35S::PIN5-GFP-3 (Mravec *et al*., 2009), PIN2::PIN2-GFP (Xu *et al*., 2006), PIN2::PIN1-GFP3 (Wisniewska *et al*., 2006), SKU5::SKU5-GFP (Sedbrook *et al*., 2002), AUX1-YFP (Swarup *et al*., 2004) and the T-DNA insertion *aux1* mutant (Alonso *et al*., 2003) were utilized. The seeds were sterilized with chlorine gas, plated on Murashige and Skoog medium (1% agar and 1% sucrose) and stratified at 4 °C for 48 hours in the dark. Seedlings were grown vertically at 21 °C under 16 h : 8 h (light : dark) photoperiod.

### DNA constructs

To fuse the green fluorescent protein (GFP) at the N-terminus of PINs, the reporter sequence, without a stop codon, was PCR amplified and connected to PIN’s start codon using the *EcoRI* (New England Biolabs) restriction enzyme. For PIN-GFP1 constructs, the GFP was inserted into PIN genomic DNA in between 45 and 46 (in PIN5) or 30 and 31 (in PIN8) amino acids, using *Xho1* and *AvrII* restriction enzymes. These PIN-GFP fusions were individually cloned into the Gateway entry vector pDONR™221 (Thermo Fishers Scientific). The 2.16 kb PIN2 promoter was cloned into the pENTR TOPO-TA vector (Thermo Fishers Scientific). The PIN2 promoter and each of the PIN-GFP entry clones were recombined into the destination vector pH7m24GW_3 (Karimi et al., 2007) by performing gateway LR reaction.

To prepare the PIN C-terminus GFP fusions, the PIN genomic DNA without a stop codon (−6 to 1889 bp for PIN5 and -6 to 1776 bp for PIN8) was cloned into the pDONR™221 vector. Next, these entry clones were individually recombined with the PIN2 promoter and pDONR™ P2r-P3 (GFP entry clone), and the resulting expression clones were recombined into the destination vector pK7m34GW (Karimi *et al*., 2005). Transgenic *Arabidopsis thaliana* plants (Columbia ecotype) were generated by performing a floral dip using the *Agrobacterium tumefaciens* (strain GV3101). List of primers used in the study is available in the supplementary information (Table S1).

### The primary root length observation

To check the functional activity of the above described PIN-GFP fusions, we observed the primary root phenotype in the PIN-GFP expressing *Arabidopsis thaliana* transgenic lines. The seedlings were grown vertically on standard MS+ media for 6 to 8 days. The seedlings were scanned, and the primary root length was measured using the ImageJ software (https://imagej.nih.gov/ij/).

### Immunocytochemistry

To map the membrane topology of PIN5 and PIN8, we implemented the whole-mount in situ immunodetection according to the previously described protocol (Gidda *et al*., 2009; Pasternak *et al*., 2015; Nodzyński *et al*., 2016). We used the anti-GFP or anti-BiP primary antibodies (1:500, Sigma-Aldrich) raised in Mouse, and the anti-Mouse secondary antibody conjugated with CY3 (1:600, Sigma-Aldrich).

### Acidification or alkalization treatment

To verify the topology of PINs, we performed selective acidification or alkalization of the apoplast using the protocol described previously in Nodzyński *et al*., (2016). To map the topology of ER PINs in reference to the ER membrane, the cytosolic pH was lowered by HCl (pH 5.0) co-treatment with digitonin (10 μM) for 30 minutes.

### Confocal Microscopy and Fluorescent Signal Analysis

We observed the GFP, RFP, anti-GFP, and anti-BiP fluorescent signals using the confocal microscopy (Carl Zeiss LSM 700 or 780 system). All images were taken with a 40X water objective. The fluorescent signal was quantified using the Fiji software (https://fiji.sc). Figures were assembled in Inkscape (inkscape.org).

## Results

To determine the membrane topology of PIN5 and PIN8 proteins, in terms of the sub-cellular orientation of their hydrophilic loop (HL) and the two terminals, we utilized various PIN-GFP fusion proteins expressed in *Arabidopsis thaliana* root cells. The previously published 35S::PIN8-GFP (Ding *et al*., 2012), and PIN2::PIN5-GFP constructs (Ganguly *et al*., 2014), which contain the GFP tag in their central hydrophilic loop (HL) allowed us to map the sub-cellular orientation of the HL. To map the sub-cellular position of the two terminals of both proteins, we generated the PIN2::PIN5-GFP and PIN2::PIN8-GFP fusions, which contain the GFP tag at their N- or C-terminal ends.

### PIN5 and PIN8 localize at the endoplasmic reticulum

To check whether the N- or C-terminal GFP fusions of PIN5 and PIN8 localize at the endoplasmic reticulum (ER), we performed a Brefeldin A (BFA) treatment, and immuno-localization utilizing the anti-BiP antibody, where BiP is an ER-localized chaperone (Gething, 1999). If these PIN-GFP fusions localize at the ER, the GFP signal should co-localize with the anti-BiP immuno-staining, and should not show BFA induced intracellular aggregations. A clear co-localization of the PIN-GFP fusions with the BiP ER marker (Figures S1a, b), and the absence of BFA aggregations (Figure S1c), indicate that these PIN-GFP fusions, similar to the previously described PIN5 (Mravec *et al*., 2009) and PIN8 proteins (Ding *et al*., 2012), localize to the ER.

### PIN5 and PIN8 exhibit antagonistic effect on the primary root growth

The previous study indicated that PIN8 promotes hypocotyl elongation while PIN5 does the inverse (Ding *et al*., 2012). In addition, Ganguly *et al*., (2014) showed that the PM localized PIN5 also inhibits primary root growth. Therefore, to assess the functionality of our newly generated PIN-GFP fusions, we investigated the PIN-mediated primary root inhibition or elongation. In comparison to the wild type Col-0, both the N- and C-terminal GFP fusions of PIN5 exhibit a significantly shorter primary root (Figure 1a). In contrast, a significantly longer roots were observed in the PIN8-GFP fusions (Figure 1b). These findings agree with the previously reported primary root phenotype of PIN8-GFP overexpression line (Ding *et al*., 2012) (Figure 1c) and PIN5-GFP fusion (Ganguly *et al*., 2014) (Figure 1d), showing that all our newly generated lines are functional. Similar to the above shown PIN5-GFP fusions, the previously published PIN5-GFP-3 overexpression line (Mravec *et al*., 2009) also inhibited the primary root growth (Figure 1c). Furthermore, the auxin influx carrier AUX1-YFP also inhibited the primary root growth while the T-DNA insertion *aux1* mutant exhibited a longer primary root (Figure 1d), indicating that auxin influx has an inhibitory effect on primary root growth. Therefore, the differences in the primary root growth in the PIN5- and PIN8-expressing *Arabidopsis thaliana* transgenic lines could be related to their opposite effects on the intracellular auxin level.

**Figure 1.**
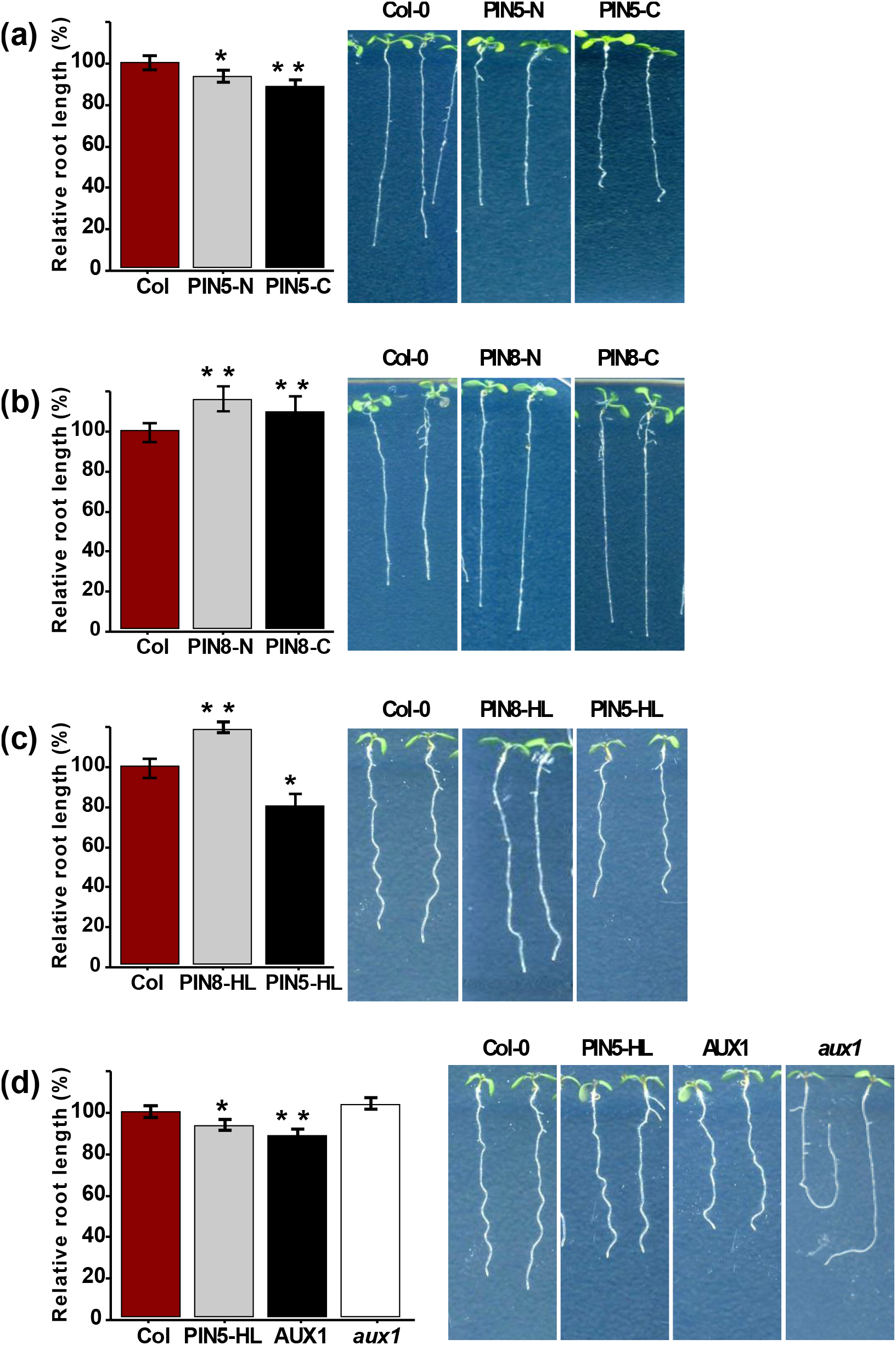
PIN5 and PIN8 act antagonistically on primary root growth. (a) PIN5 N- and C-terminal GFP fusions expressed under PIN2 promoter. (b) PIN8 N- and C-terminal GFP fusions expressed under PIN2 promoter. (c) PIN8-GFP and PIN5-GFP-3 overexpression lines which contain the GFP in their HL and expressed under 35S promoter. (d) PIN5 protein which contains GFP in its HL and expressed under PIN2 promoter, AUX1 protein which contains the N-terminal yellow fluorescent (YFP) tag expressed under its native promoter. The wild type (Col-0) was used as a control and its root length is calculated to 100%. The bar graphs indicate relative root length in comparison to Col-0. The root pictures next to each bar are representative images for the root length. The error bars represent SEM. The asterisks indicate significant differences in comparison to WT (Col-0) (* P<0.05, ** P < 0.01, Student’s t-test).

### The N-terminal end of PIN5 faces the cytoplasm while its C-terminus resides in the ER lumen

The functionality of the PIN-GFP fusions mentioned above allowed us to use them as valid tools for mapping their membrane topology. First, we mapped the topology of PIN5. We performed immunodetection using IGEPAL (2%) to permeate all cellular membranes (Pasternak *et al*., 2015; Nodzyński *et al*., 2016) or digitonin (40 μM) to selectively permeabilize the PM alone (Lorenz *et al*., 2006; Gidda *et al*., 2009). Permeabilization with IGEPAL enables the labelling of epitopes situated both in the cytoplasm and in the ER lumen (Gidda *et al*., 2009). However, permeating the plasma membrane with digitonin enables to label the epitopes residing in the cytosol but not inside the ER. Simultaneous immunodetection using the two protocols enables to differentiate between an epitope positioned in the cytoplasm or in the ER lumen.

In a control experiment carried out to label the plasma membrane localized PIN2-GFP, which has the GFP insertion in its cytoplasmic central hydrophilic loop, the antibody detected the GFP both in the IGEPAL and digitonin permeated cells (Figures 2b, c). In addition, the quantified anti-GFP signal after permeabilizing the cells with either IGEPAL or digitonin was similar (Figure 2j), indicating that digitonin permeates the plasma membrane as effectively as IGEPAL. To test whether the concentration of digitonin used in this experiment permeabilized the plasma membrane alone, but not the ER membrane, we performed similar experiment by using the BiP protein, a HSP70 chaperone located in the ER lumen (Gething, 1999). Permeabilization with IGEPAL allowed the immuno detection of the chaperone (Figure 2d, k). However, after permeating the cell by digitonin, the anti-BiP antibody did not label the protein (Figures 2e, k), showing that the antibody did not access the luminal BiP because the ER membrane remained intact.

**Figure 2.**
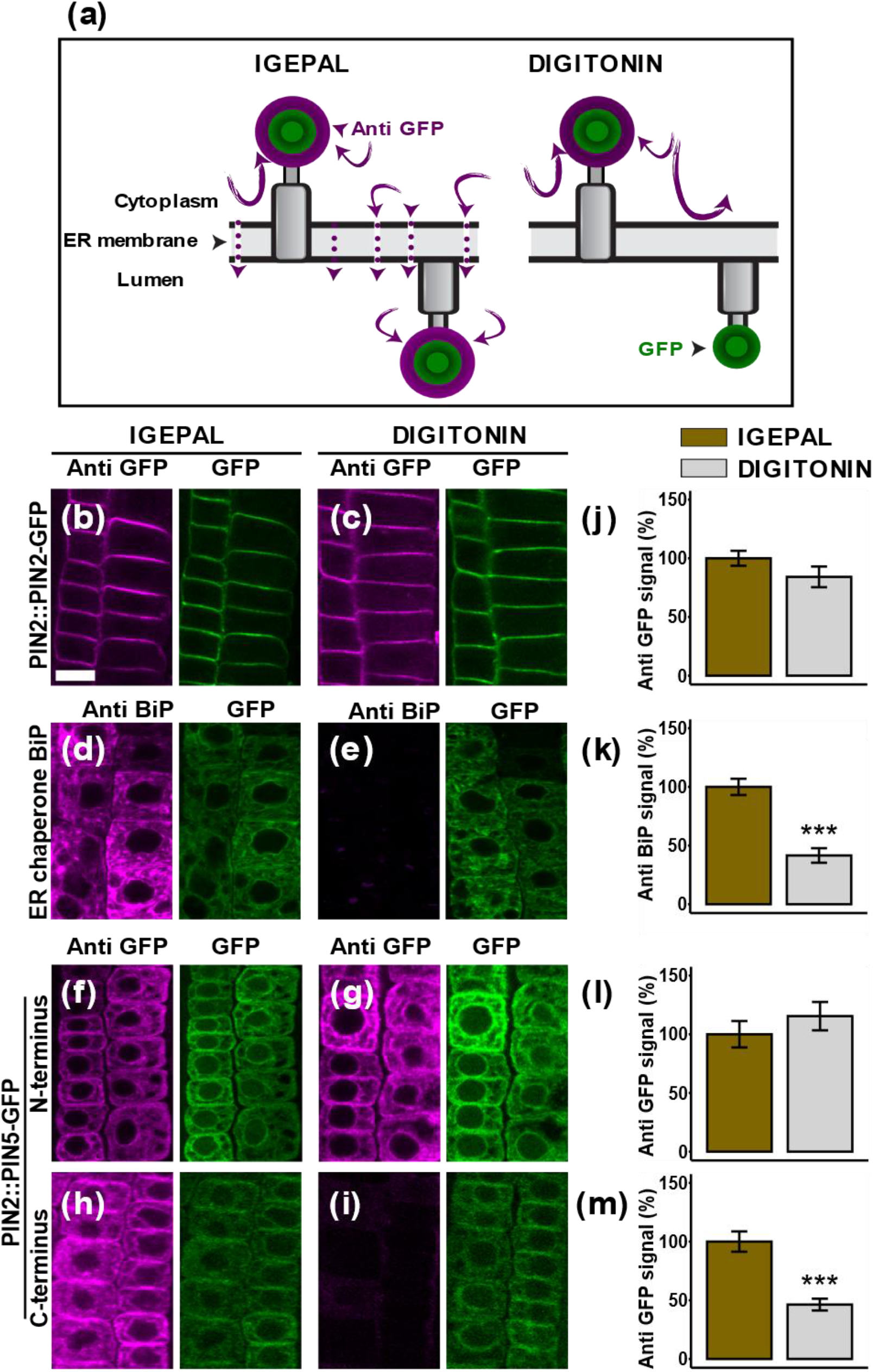
PIN5 has its N-terminus in the cytoplasm and its C-terminus in the ER lumen. (a) Cartoons showing the principle of ER membrane permeable versus non-permeable immunocytochemistry. IGEPAL permeates all cellular membranes and allows antibodies to detect epitopes positioned in the cytoplasm and the ER lumen. Digitonin permeates the PM alone but not the ER membrane and enables to label the cytoplasmic epitope alone, but not the luminal epitope. (b-i) Representative epidermal root cell pictures for immunolocalization. The designations anti-GFP or GFP refer to the immuno-staining and the native GFP fluorescence signal respectively while the anti-BiP refers to the BiP immuno-staining signal. To label the ER chaperone BiP, as a luminal control, PIN2::PIN5-GFP construct was used. (j-m) Quantified immunofluorescence signal intensity. The IGEPAL (control) immuno fluorescence is plotted as 100%. The asterisks indicate significant differences in comparison to PM permeabilization with IGEPAL (*** P < 0.001, Student’s t-test). The significantly lower signal after digitonin treatment compared to IGEPAL indicates the ER lumen orientation of protein moiety. The error bars represent SEM from the total number of seedlings analyzed from three biological experiments (n > 15 per experiment). Scale bar = 10μm.

Consistently with the PIN2-GFP immunodetection, the GFP tag at the N-terminal end of PIN5 was clearly detected after permeabilization with IGEPAL or digitonin (Figures 2f, g, l), indicating that the N-terminal end of the protein is oriented cytoplasmically. In addition, we performed a similar experiment to label the PIN5-GFP-1 insert. This construct contains the GFP insertion in the first small loop, which is connected to the first TM helix at the opposite side of the N-terminal end (Figure S6b). The anti-GFP labelled the GFP-1 moiety only in the IGEPAL permeated cells (Figures S3a, b, e). This result implies that the first minor loop which is in between the first two N-terminal helices of PIN5 is facing the ER lumen. This data also supports the finding which indicate that the amine end of this protein is in the cytoplasm. However, the C-terminal end of PIN5-GFP was detected in the IGEPAL permeated cells alone, but not in the digitonin permeated cells (Figures 2h, i, m), showing that the carboxy terminus of PIN5 may be oriented on the luminal side of the ER membrane.

### The hydrophilic loop of PIN5 localizes in the cytoplasm

Although PIN5 localizes at the ER (Mravec *et al*., 2009), the plasma membrane localization of this protein (Ganguly *et al*., 2014) provided the possibility to determine the topology of its hydrophilic loop (HL), in which the GFP was inserted. We performed immunocytochemistry with and without permeating the plasma membrane. In this study, the immuno-localization technique in which the plasma membrane was permeated by IGEPAL after tissue fixation with paraformaldehyde (PFA) (Pasternak *et al*., 2015) and glutaraldehyde (GA) is referred to as the plasma membrane permeable protocol. This technique allows the labelling of an intracellular and an apoplastic epitope. Excluding membrane permeabilization detergents from the protocol enable to maintain the intact plasma membrane. This technique is limited to label extracellularly positioned epitopes alone and is referred to as the membrane non-permeable protocol (Nodzyński *et al*., 2016).

In a control experiment carried out using the extracellularly positioned SKU5-GFP, the epitope was clearly labelled both in the membrane permeable and non-permeable immuno detections (Figures 3b, c, j). However, the anti-GFP antibody labelled the cytoplasmic PIN1-GFP3 (Figures 3d, e, k) only upon membrane permeabilization with IGEPAL. These results indicate that the protocol was effective enough to distinguish between the apoplastic and the cytoplasmic epitopes. Similarly, PIN5-GFP was clearly labelled under the membrane permeable immunolocalization alone, while detection of the protein was abolished in the membrane non-permeable protocol (Figures 3f, g, l), indicating that the HL of PIN5 localizes in the cytoplasm. Moreover, in an analogical experiment conducted with PIN5 (C1-GFP), PIN5 chimeric protein which contains GFP tagged PIN2-HL in the PIN5-HL (Ganguly *et al*., 2014), the antibodies label the C1-GFP moiety in the membrane permeable immunolocalization alone (Figures 3h, i, m). Therefore, if the GFP epitope cannot be detected without membrane penetration, this strongly supports that the hydrophilic loop of the PIN5 is found in the cytoplasm.

**Figure 3.**
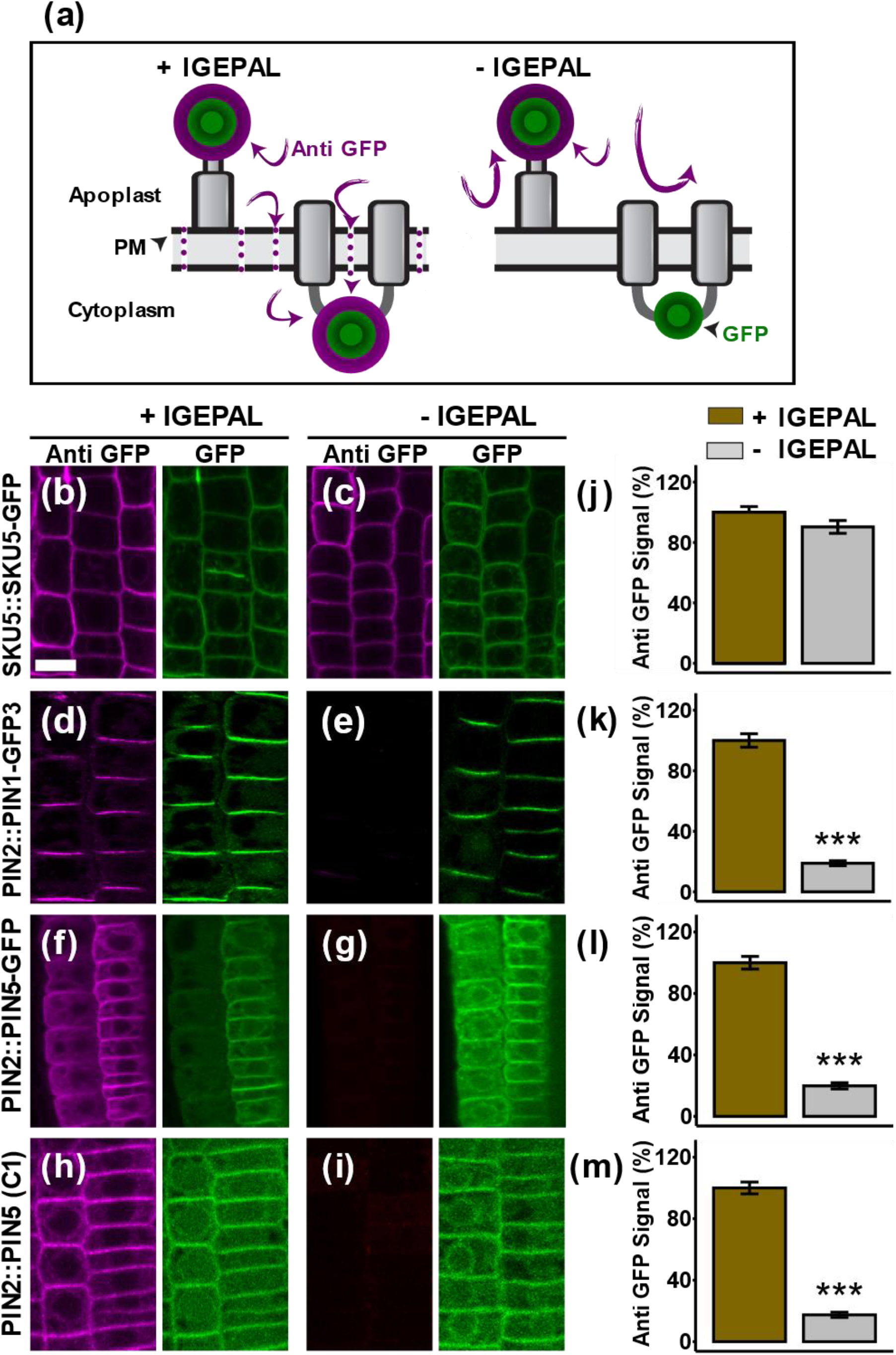
The central hydrophilic loop of PIN5 is found in the cytoplasm. (a) Cartoon showing the principle of PM permeable vs. non-permeable immunostaining. PM permeable protocol using IGEPAL allows to label both apoplastic and cytoplasmic epitopes. The PM non-permeable protocol in the absence of IGEPAL limited the antibody to detect only the apoplastic epitope. (b-i) Representative root pictures for PM permeable and PM non-permeable immunolocalization. The plasma membrane permeable (+ IGEPAL) immuno-fluorescence is plotted as 100%. The asterisks indicate significant differences in comparison to PM permeabilization with IGEPAL (*** P < 0.001, Student’s t-test). The presence or absence of anti-GFP signal in the plasma membrane non-permeable condition (−IGEPAL) indicates the apoplastic or cytoplasmic orientation of the epitopes respectively. (j-m) Quantified immunofluorescence signal intensity. Error bars represent SEM from the total number of seedlings analyzed from three biological experiments (n > 18 per experiment).

### The N- and C-terminal ends of PIN8 are oriented towards the cytoplasm while its hydrophilic loop faces the ER lumen

Next, we mapped the topology of PIN8. To test more controls in these sets of experiments, we used the cytoplasmic PIN1-GFP3 (Nodzyński *et al*., 2016). As expected, like the PIN2-GFP (Figures 2b, c, j), the GFP3 moiety was clearly detected after permeating the cell with either IGEPAL or digitonin (Figures 4a, b, k). However, the ER-luminal BiP immuno detection revealed almost no signal in digitonin-permeated cells in comparison to the IGEPAL treatment (Figures 4c, d, l). This consistently demonstrated that the ER lumen is not accessible to antibodies when digitonin alone is used in the immuno-localization protocol. In the case of the PIN8-GFP fusions, similar to the PIN1-GFP3, the N-terminal (Figures 4e, f) and the C-terminal (Figures 4i, j) were clearly labelled with the anti-GFP antibody both in the IGEPAL and digitonin-permeated cells. There was statistically no significant difference between the quantified anti-GFP signal corresponding to IGEPAL and digitonin treatment both in the PIN1-GFP3 (Figure 4k) and PIN8-GFP N- and C-terminus (Figures 4m, o). These results indicate that the two terminal ends of PIN8 protein orient on the cytoplasmic side of the ER membrane. In contrast, the GFP immuno detection of PIN8-GFP1, in which we inserted the GFP in between the first two helices of the N-terminal domain (Figure S6b), was significantly lower in digitonin permeated cells, in comparison to the IGEPAL permeabilized cells (Figures S3c, d, f). This result shows that PIN8 contains its first minor loop in the ER lumen, and this conclusion supports the cytoplasmic orientation of its N-terminus. Furthermore, to determine the orientation of PIN8 HL, we took the advantage of the previously generated ER-localized 35S::PIN8-GFP (Ding *et al*., 2012). The immuno detection of the central HL of PIN8 showed markedly weaker labelling in digitonin permeated cells in contrast to permeabilization with IGEPAL (Figures 4g, h, n). This shows that PIN8 contains its central HL in the ER lumen, which is oriented to the opposite position of its terminus.

**Figure 4.**
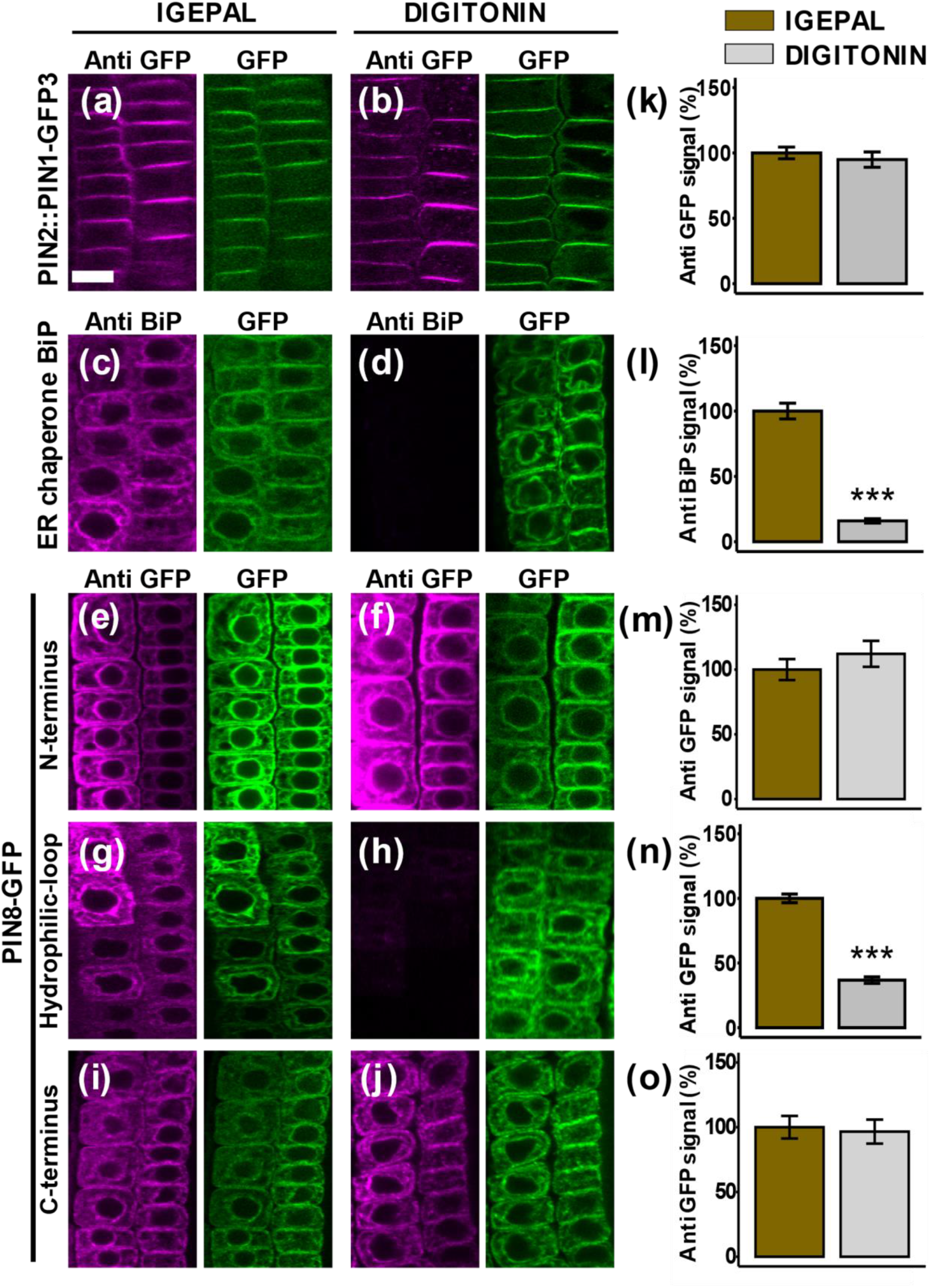
PIN8 has its N - and C - terminus in the cytoplasm while its central hydrophilic loop localizes in the ER lumen. (a-j) Representative root pictures after anti-GFP immunolocalization. The anti-GFP or GFP labels in the figures indicate the anti-GFP or GFP fluorescence signal while anti BiP is the anti BiP immunodetection fluorescent signal. To label the ER chaperon BiP as a luminal control, PIN2::PIN8-GFP construct was used. (k-o) Quantified immunofluorescence signal intensity. The immunofluorescence after the IGEPAL permeabilization (control) is plotted as 100%. The asterisks indicate significant differences in comparison to permeabilization with IGEPAL (*** P < 0.001, Student’s t-test). The significantly lower immuno-fluorescence signal after digitonin treatment compared to the IGEPAL indicates the ER-luminal orientation of the reporter. Error bars represent SEM from total number of seedlings analyzed from three biological experiments (n > 15 per experiment). Scale bar = 10μm.

### Selective acidification of the cytoplasm or the apoplast largely corroborates the membrane topology of PIN5 and PIN8

To verify the results described above, we utilized GFP as a pH-sensitive probe (dos Santos *et al*., 2020), and adopted a fluorescent protein quenching assay (Swarup et al., 2004; Nodzyński *et al*., 2016) for studying the topology of ER-localized PIN-GFP proteins in a live root cell. To evaluate the applicability of the method, we first assessed the effect of the HCl (pH = 5.0) on the fluorescent reporter signal. The PIN2-GFP in which the fluorescent moiety faces the cytoplasm, the HDEL-RFP which resides in the ER lumen, and PIN5-GFP and PIN8-GFP fusion proteins which localize at the ER were incubated (30 minutes) in liquid Murashige and Skoog medium (MS, pH = 5.9 (control)) or the MS medium titrated with hydrochloric acid (HCl, pH = 5.0). The HCl should not affect the GFP and RFP fluorescence of the intracellular chimeric proteins because the PM is not permeable to the protons generated after HCl dissociation in the titrated MS medium (Nodzyński *et al*., 2016). Both in the PIN2-GFP and HDEL-RFP, the acid treatment did not affect the fluorescent signal (Figures 5b, c, g, h). Similarly, the HCl did not quench PIN5-GFP and PIN8-GFP fusion proteins (Figures 5d-f, i-k). These results indicate that acidification of the apoplast does not affect the intracellular reporters. Therefore, to utilize the HCl-mediated fluorescent quenching for studying the topology of a protein in reference to the ER membrane, the PM needs to be permeated.

**Figure 5.**
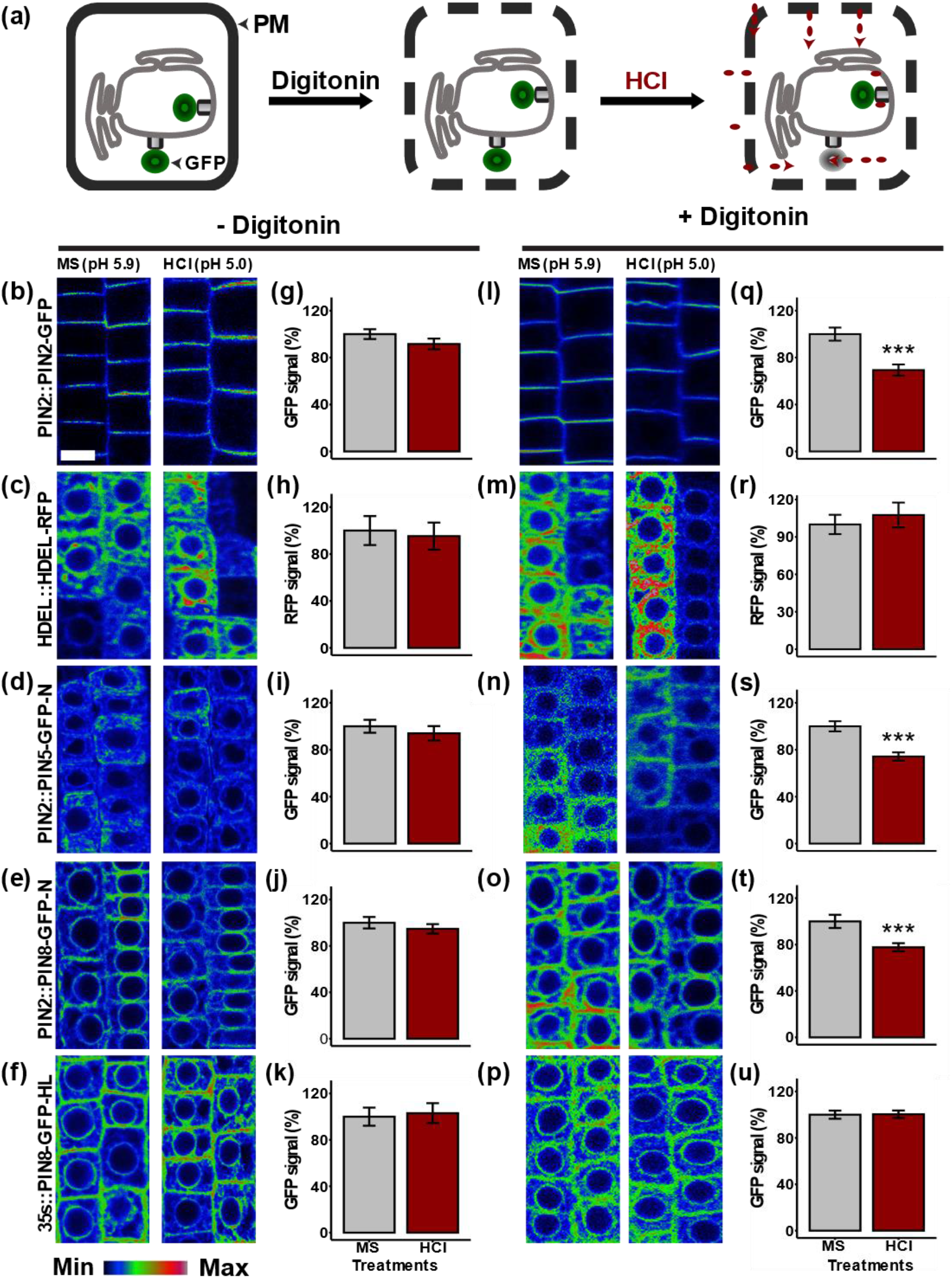
In the absence of digitonin, the PM non-permeable HCl does not quench the intracellular fluorescent while permeating the PM with digitonin allows quenching the cytoplasmic fluorescent. (a) Cartoon showing the principle of the intracellular GFP quenching. Acidification of the cytosol after permeating the PM with digitonin decreases the fluorescence of the cytoplasmic GFP reporters while the ER-luminal ones are not affected. (b-f) Color-coded root pictures from a live cell imaging show that acidification of the apoplast did not affect the intracellular fluorescent reporter signal. (g-k) Quantified fluorescent signal from seedlings subjected to acidification of the apoplast. (l-p) Pictures from root cells exposed to acidification of the cytoplasm. (q-u) Quantified fluorescent signal after acidification of the cytoplasm. The GFP signal in the MS (control) treatment is plotted as 100%. The asterisks indicate significant differences in comparison to the control MS treatment (*** P < 0.001, Student’s t-test). Error bars represent SEM from the total number of seedlings analyzed from three biological experiments (n > 15 per single experiment). Scale bar = 10μm.

To permeate the PM in live root cells, we optimized the minimum concentration of digitonin (10 μM), which should not affect the PIN-GFP fluorescent signal. To assess the effect of digitonin on the GFP signal, the cytoplasmic PIN2-GFP, and the ER localized PIN8-GFP and PIN5-GFP transgenic lines were incubated in liquid MS medium with and without digitonin. The GFP signal in all constructs subjected to digitonin treatment was similar to the lines incubated in the control MS medium without digitonin (Figure S2). These results show that the concentration of digitonin used to permeate the membrane in live root cells did not affect the GFP fluorescent signal.

To permeate the PM and to acidify the cytosol, six days old seedlings expressing PIN-GFP fusions were subjected to digitonin co-treatment with either MS (control) or HCl. If digitonin permeates the PM in a live root cell, but not the ER membrane, the fluorescent signal of GFP situated in the cytoplasm should be decreased in response to the acid treatment while the one which is enclosed in the ER lumen should remain unaffected. After permeating the PM with digitonin, the acid treatment quenched the cytoplasmic PIN2-GFP and significantly decreased the fluorescent signal in comparison to the respective MS control treatment (Figures 5l, q). This result indicates that digitonin perforates the PM which allows the HCl to diffuse into the cytoplasm and quenches the GFP moiety. To check whether digitonin permeates only the PM, but not the ER membrane, an analogical experiment was conducted by using HDEL-RFP, in which the RFP moiety is enclosed in the ER lumen. If the digitonin permeates the ER membrane, the HDEL-RFP signal should be diminished in response to the acid treatment. However, after the digitonin and acid co-treatment, the RFP signal remained as high as the fluorescent signal in the control treatment (Figures 5m, r). To check whether the HDEL-RFP stability in this experiment was due to the RFP resistance to the lower pH or because it was protected by the ER membrane, we performed similar experiment by using MS medium titrated with propionic acid (pH = 5.0). Since the propionic acid is membrane permeable, it should quench the luminal fluorescent. As expected, the HDEL-RFP signal was significantly degraded after treatment with the propionic acid (Figure S4). These results collectively suggest that the insensitivity of the RFP to the HCl, although the PM was permeated with the digitonin, is due to its enclosure in the ER lumen, but not because of its resistance to the acidic pH.

After permeating the PM with digitonin, similar to the observation in the PIN2-GFP, the fluorescent signal both in the PIN5 and PIN8 N-terminal GFP tag was significantly quenched by the HCl (Figures 5n, o, s, t), confirming that the N-termini of both proteins are in the cytoplasm. Subjecting the GFP moiety to the acidic pH titrated with HCl does not cause protein degradation (Nodzyński *et al*., 2016). The GFP signal loss at the acidic pH is related to the pH-induced secondary structural distortion of the tag (dos Santos *et al*., 2020). However, both the PIN5-GFP1 and PIN8-GFP1 lines, which contain the GFP in the small loop extending from the first TM-helices at the opposite side of the N-terminus, remained stable after the digitonin and acid co-treatment (Figures S3 g-j). These results indicate that the first minor loop in the N-terminal domain of both PIN5 and PIN8 is in the ER lumen. Similarly, the GFP moiety in the PIN8 hydrophilic loop was not affected by the acidification of the cytosol (Figures 5p, u), showing that the HL of the protein is enclosed in the ER lumen. These results agree with the immunocytochemistry findings.

Furthermore, to check the sub-cellular localization of PIN5 HL, we took the advantage of the ectopic PM localization of PIN2::PIN5-GFP line, which contains the GFP insertion in its central HL (Ganguly *et al*., 2014). This transgenic line was subjected to either acid or base treatment using MS (control, pH = 5.9) titrated with HCl (pH = 5.0) or buffered with KOH (pH = 8.0). Both HCl and KOH are PM non-permeable (Nodzyński *et al*., 2016). If the reporter moiety is extracellularly localized, the acidic pH decreases the fluorescent signal while the alkaline treatment enhances it (Swarup *et al*., 2004; Nodzyński *et al*., 2016). The glycosyl phosphatidylinositol-anchored SKU5-GFP, in which the fluorescent reporter is positioned at the exterior of the cell (Sedbrook *et al*., 2002) and the intracellular PIN2-GFP were used as control constructs with a known topology.

The acid treatment significantly decreased the fluorescent signal in the apoplastic SKU5-GFP (Figures 6b, c, n) in comparison to the respective control medium (MS, pH = 5.9), while a non-significant change was observed in PIN2-GFP (Figures 6e, f, o) and PIN5-GFP (Figures 6h, i, p). In addition, we performed similar experiment by utilizing the C1 construct (PIN5-PIN2-HL-GFP) to further verify the orientation of PIN5 HL. Like our observation in the PIN2-GFP and PIN5-GFP, the acid treatment did not affect the GFP signal in the C1 (Figures 6k, l, q). Furthermore, the alkalization treatment (KOH, pH = 8.0) enhanced the GFP signal only in the SKU5-GFP (Figures 6b, d, n), but not in the other transgenic lines (Figures 6g, j, m, o, p, q). These results support the above mentioned immunolocalization findings (Figures 3f, g, h, i, l, m), which show that the HL of PIN5, like that of PIN2, is oriented in the cytoplasm.

**Figure 6.**
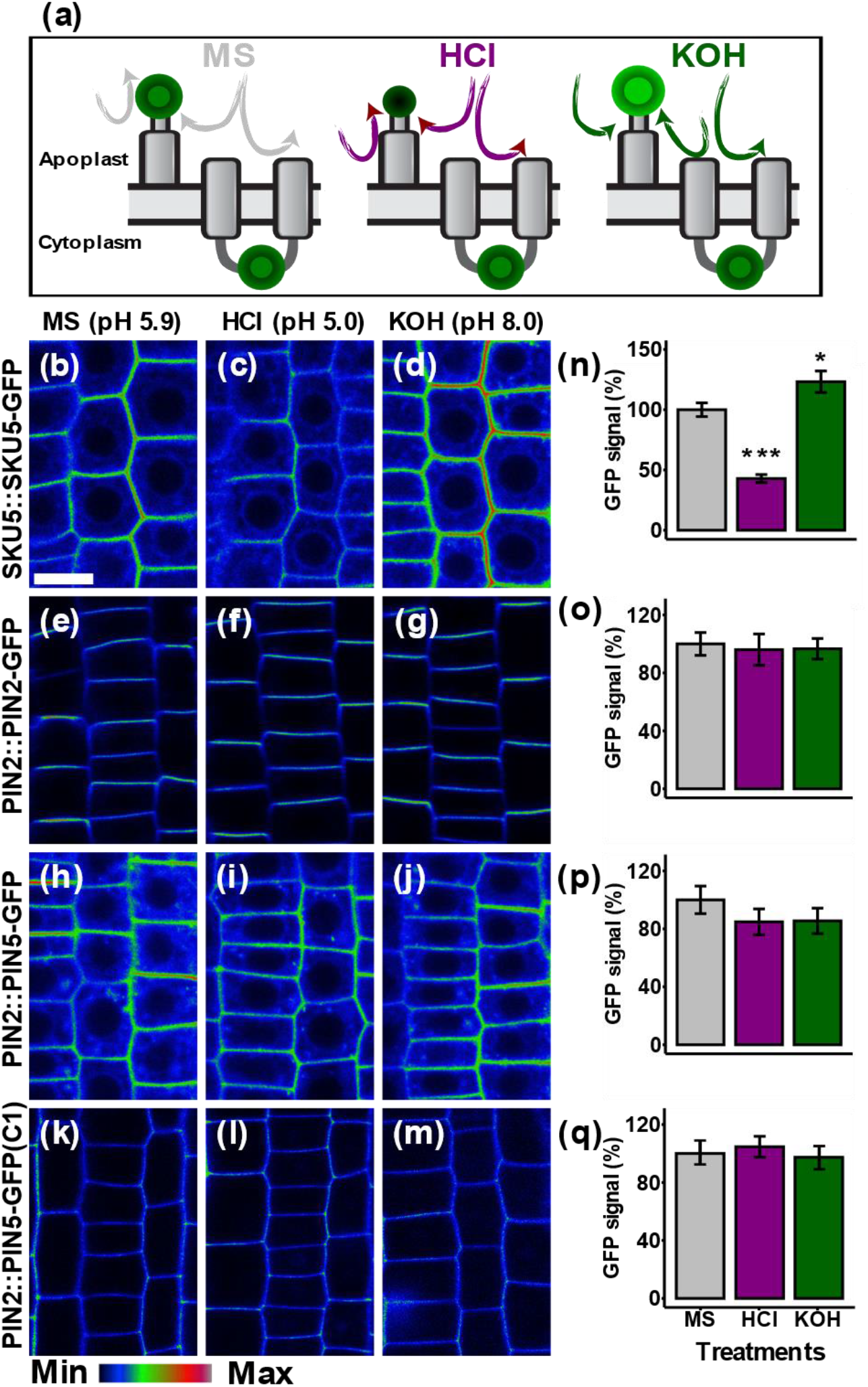
GFP quenching reveals the cytoplasmic position of the PIN5 hydrophilic loop. Six days old seedlings were subjected to standard Murashige Skoog Medium (pH = 5.9, control) buffered with either HCl (pH = 5.0) or KOH (pH = 8.0) for 30 minutes. (a) Cartoon showing the principle of the extracellular GFP quenching. (b-m) Root images are shown in signal intensity color code to better visualize the GFP intensity changes after the acidic or alkaline treatment. (n-q) Quantified GFP signal intensity. The control treatment (MS) per each construct was plotted as 100%. The asterisks indicate significant differences in comparison to the control MS treatment (*** P < 0.001, *P < 0.05, Student’s t-test). The error bars represent SEM from the total number of seedlings obtained from three biological experiments (n > 18 per single experiment).

## Discussion

Although the mutually opposing role of PIN5 and PIN8 proteins in regulating the intracellular auxin homeostasis (Mravec *et al*., 2009; Ding *et al*., 2012), and various developmental aspects (Ganguly *et al*., 2010; Ding *et al*., 2012; Sawchuk *et al*., 2013; Lee *et al*., 2020) have been reported, how the two proteins which localize at the same intracellular compartment (endoplasmic reticulum) antagonize each other remains unclear (Ding *et al*., 2012). In this study, we determined the membrane topology of PIN5 and PIN8, in terms of the sub-cellular orientation of their N- and C-terminal ends, and the central hydrophilic loop. Our results showed that, despite the similarities in the orientation of the N-terminus, PIN5 and PIN8 have divergent membrane topologies. Therefore, the mutually opposing activity of the two proteins could be attributed to their topological differences.

The cytoplasmic and the ER luminal localization of the N- and C-terminus of PIN5 respectively (Figure 2), implies that the protein’s topology is composed of an odd number of transmembrane domains. In addition to this, the luminal position of the first small loop in the N-terminal domain (Figures S3) and the cytoplasmic position of its central hydrophilic loop (Figure 3) indicate that PIN5 has four and five transmembrane helices in its N- and C-terminal domain, respectively. These PIN5 structural data do not agree with the 3D structure of the AlphaFold model (Jumper *et al*., 2021). The quality of the AlphaFold structural model is determined by a per-residue confidence score (pLDDT) which ranges between 0 and 100. The pLDDT for the prediction of PIN5 terminal ends and the central HL varies from low (70 > pLDDT > 50) to very low (pLDDT < 50) score (Figure S6a). The authors declared that regions with a very low pLDDT might be unstructured in isolation. In fact, most structural models are prone to this limitation. For example, ARAMEMNOM predicted 10 TMDs of the AUX1 protein, however, an experimental topology study with AUX1-YFP quenching demonstrated that this protein has 11 TMDs (Swarup *et al*., 2004). In the study, the algorithm failed to differentiate the boundary between the AUX1 TM helices. Although AlphaFold often provides almost an accurate prediction of transmembrane protein structure, it does not consider the essential post-translational modifications which affect the structure and function of proteins (Bagdonas *et al*., 2021). As a result, AlphaFold 3D structural model may not always resemble the native topology of a protein (Azzaz & Fantini, 2022). Hence, the discrepancy between the experimentally verified topology of PIN5 versus the predicted one could be due to the limitation of the algorithms to correctly identify the unstructured hydrophilic loop. Therefore, it is likely that PIN5 has 9 TMDs topology, with the central HL positioned after the 4^th^ alpha helix (Figure S6b).

The cytoplasmic position of both termini of PIN8 (Figure 4) indicates that this protein has an even number of transmembrane domains. This finding agrees with the topology model obtained from UniProt (Bateman *et al*., 2015), which predicted ten alpha helices of the PIN8 structure. In addition, AlphaFold databases (Jumper *et al*., 2021; Varadi *et al*., 2022) indicated a similar orientation of the two termini to the opposite position of the central hydrophilic loop (Figure S6a), showing that the protein has an even number of TMDs. Furthermore, the crystal structure of PIN8 also demonstrated that this protein has 10 TMDs (Ung *et al*., 2022). However, in contrast to our findings, in terms of the ins and outs orientation of the two termini and the central hydrophilic loop of the protein, Ung *et al*., (2022) reported that the hydrophilic loop of PIN8 localizes in the cytoplasm while the two termini possess non-cytoplasmic position. The reason for the contradicting position of the PIN8 structure could be related to the method utilized. In our study, we mapped the membrane topology of the protein in *Arabidopsis thaliana* root cells, while the protein is in its biological membrane. However, Ung *et al*., (2022) determined the structure of the protein with Cro-EM using purified and crystallized protein. Therefore, it might be possible that the processes of isolating, purifying, crystallizing, and embedding a protein in nano-discs with membrane patches abolish the protein’s membrane topology (Yao *et al*., 2020), which makes it challenging to correctly show its ins and outs native structure. Hence, the most plausible topology model for PIN8 would be the one which contains 10 TMDs with groups of five alpha-helices separated by the central hydrophilic loop residing in the ER lumen (Figure S6b).

The topology of polytopic membrane proteins is determined during their biogenesis at the ER membrane. At the ER, the signal recognition particle (SRP) recognizes the first TM helix as it emerges from the ribosome and targets the nascent helices to the ER membrane, where the Sec61 translocon facilitates their integration into the lipid bilayer (Spiess *et al*., 2019). Likewise, the cytosolic exposure of the N-terminus and the luminal enclosure of the first small loop in the N-terminal domain, both in PIN5 and PIN8 indicate that the synthesis of these proteins, like other ER proteins, is based on the initial insertion of their first TM helices into the ER membrane. Once the protein is anchored in the ER membrane, its ins and outs topological orientation in terms of the cytoplasmic and the luminal domains are highly determined by the positive inside rule (VonHeijne, 1989). Based on this paradigm, although additional factors influence the protein’s structure, the positively charged residues remain in the cytoplasm while the luminal domains retain in the lumen and the TMDs embed in the lipid bilayer (Helfand *et al*., 2003; Spiess *et al*., 2019). Therefore, the negative residues in the PIN8’s central HL are likely to keep the loop in the ER lumen whereas the HL of PIN5 stays inside the cytoplasm regardless of the localization of the protein at the PM or the ER membrane due to its retention of the positive residues (Figure S5). The electrostatic potential of PIN5 and PIN8, which was predicted by using the 3D structure of the protein from the AlphaFold (Figure S5), could also show that the GFP insertion in the loop of these proteins did not change their native structure. This is because, according to the positive inside rule, the charge distribution of the AlphaFold model which does not contain the GFP sequence agrees with the experimental data, which shows the cytoplasmic and non-cytoplasmic position of PIN5 and PIN8 HL, respectively. Furthermore, studies have shown that the addition of a GFP tag to a protein of interest does not change the properties of the protein (Rizzuto *et al*., 1995; Lorenz *et al*., 2002; White *et al*., 2015).

The divergent topology of PIN5 and PIN8 may reflect their opposing activity in mediating the intracellular auxin homeostasis and developmental events. The lower level of free indole-3-acetic acid (IAA) and the increased accumulation of auxin conjugates after induction of PIN5 expression has been reported (Mravec *et al*., 2009). In contrast to PIN5, the PIN8 overexpression line exhibited a higher level of free IAA than its conjugates (Mravec *et al*., 2009; Ding *et al*., 2012). Furthermore, Ganguly *et al*., (2010) have reported that PIN5 enhances NAA accumulation in BY-2 tobacco cells, in contrast to PIN8. The authors also demonstrated that PIN5 promotes root hair growth while PIN8 inhibits it. Similarly in this study, we observed that PIN5 inhibits primary root growth while PIN8 promotes the root growth (Figure 1). Indeed, the primary root growth is inversely proportional to the cellular auxin level, and the TIR1/AFB-Aux/IAA signalling pathway is needed for auxin-induced root growth inhibition (Fendrych *et al*., 2018). Hence, the contradicting activity of PIN5 and PIN8 proteins may be related to their differential auxin transport activity across the ER membrane. The previous studies have proposed that PIN5 transports auxin into the ER lumen (Mravec *et al*., 2009), while PIN8 effluxes it out of the ER (Ding *et al*., 2012). In agreement with these hypotheses, it is possible that the cytoplasmic hydrophilic loop of PIN5 conveys auxin flux into the ER lumen, which then promotes auxin supply to the nucleus and inhibits the root growth. On the contrary, PIN8 protein, which contains its hydrophilic loop in the ER lumen, may export auxin out of the lumen, which leads to the nuclear auxin depletion and therefore prevents inhibition of root growth.

In conclusion, this study shows that PIN5 and PIN8 have overall opposite membrane topology, except for their similarity in the orientation of the N-terminal end. This points to the potential role of the topological differences between the two proteins to establish their antagonistic functionality.

## Acknowledgements

This work was supported by the Ministry of Education, Youth and Sports of the Czech Republic under the project CEITEC 2020 (project no. LQ1601). We acknowledge the Core Facility CELLIM of CEITEC supported by the Czech-BioImaging large RI project (LM2018129 funded by MEYS CR) part of the Euro-BioImaging (www.eurobioimaging.eu) ALM and medical imaging Node (Brno, CZ) Plant Sciences Core Facility of CEITEC Masaryk University for the technical support. We thank Hélène S. Robert, Jiří Friml, and Jan Hejátko for their valuable comments and ideas enhancing the experimental design of the work. We also thank Sravan Thulakumar for technical advice on molecular cloning, Surendra Saddala for technical advice in *Agrobacterium tumefaciens* transformation, Veronika Bilanovičová for valuable discussions improving the presentation clarity of the results.

## Author contributions

YS and TN designed the experiments. YS generated PIN-GFP fusions and conducted immunocytochemistry and GFP quenching experiments. YS, NR and VP performed primary root length experiment, YS and MZ tested the effect of digitonin on GFP signal, YS and TN wrote the manuscript.

## Data availability

The data that support this study and all generated constructs including plasmids and transgenic lines will be made available from the corresponding author (Tomasz Nodzyński) up on request.

