## Supplemental figures for "The intracellular auxin homeostasis regulators PIN5 and PIN8 have a divergent membrane topology in *Arabidopsis thaliana* root cells"

### Supplementary information

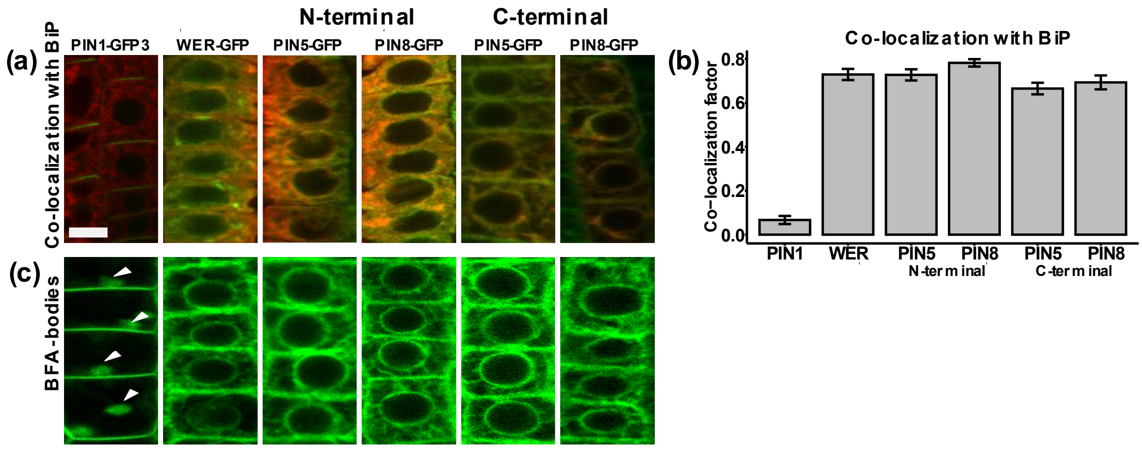


**Figure S1.** PIN5 and PIN8 localize to the ER**.**

The N- and C-terminally GFP tagged PIN5 and PIN8 localize at the ER. The PIN-GFP fusions were expressed under PIN2 promoter in the *Arabidopsis thaliana* root cell. PIN1-GFP3 and WER-GFP were utilized as a control for the plasmamembrane and endoplasmic reticulum localization respectively. (a-b) PIN5 and PIN8 co-localize with the ER chaperon BiP. The co-localization factor was determined using Zeiss software, where the co-localization factor close to 1 shows strong co-localization. (c) The BFA aggregation in the plasma membrane localized PIN1-GFP3 (indicated by the white arrows) and the absence of BFA aggregations both in PIN5 and PIN8 indicates the absence of these proteins from the PM derived endosomes. Five days old seedlings were incubated in BFA (25 µM) for one hour. For microscopy observation, 5 days old seedlings (n = 5) and at least 10 epidermal root cells per seedling were included in the analysis. The error bars represent the SEM. Scale bar, 10 µm.

### Supplementary information

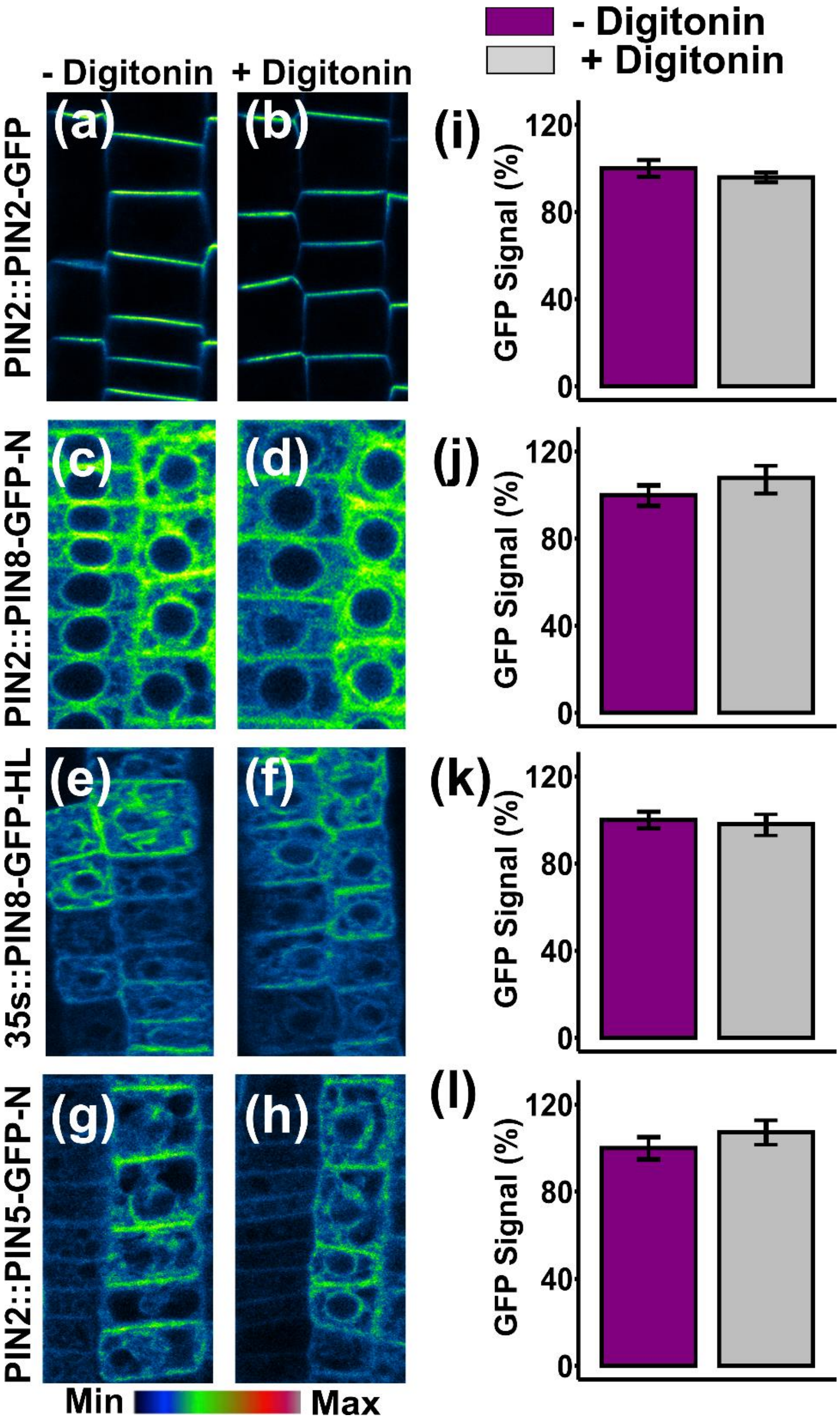


**Figure S2.** Digitonin does not affect the green fluorescent protein signal.

(a-h) Epidermal root cell images shown in signal intensity color code to better visualize the GFP intensity changes before and after digitonin treatment. Six days old seedlings were treated by either MS+ (control) or MS+ with digitonin for 30 minutes. (i-l) Quantified GFP fluorescent signal. The error bars represent SEM from three independent biological experiments (n > 10 per experiment). Scale bar, 10 µm.

### Supplementary information

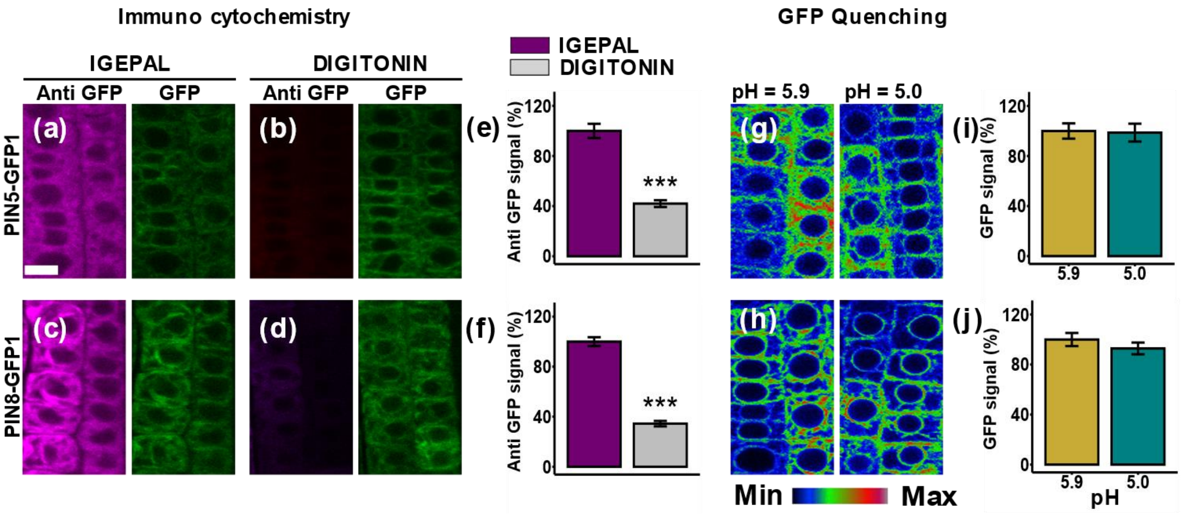


**Figure S3.** Immuno-cytochemistry and GFP quenching of the PIN-GFP1 insert.

The GFP was inserted in between the first and the second predicted helices both in PIN5 and PIN8. (a-d) The immuno-detection of the GFP is abolished in digitonin permeated epidermal root cells. (e-f) The quantified anti GFP signal. The control treatment (IGEPAL) was plotted as 100%. The asterisks indicate significant differences in comparison to PM permeabilization with IGEPAL (*** $P < 0.001$, Student's t-test). The significantly lower immuno-fluorescence signal after digitonin treatment compared to the IGEPAL indicates the ER-luminal orientation of the reporter. The error bars represent SEM from the total number of seedlings obtained from three biological experiments (n > 18 per single experiment). (g-h) Root epidermal cell images shown in signal intensity color code to better visualize the GFP intensity changes after the acidic or basic treatment. Six days old seedlings were co-treated by either digitonin and MS+ or digitonin and HCl for 30 minutes. (i-j) Quantified GFP fluorescent signal. Error bars represent SEM from the total number of seedlings analysed from three biological experiments (n > 15 per single experiment). Scale bar, 10 µm.

### Supplementary information

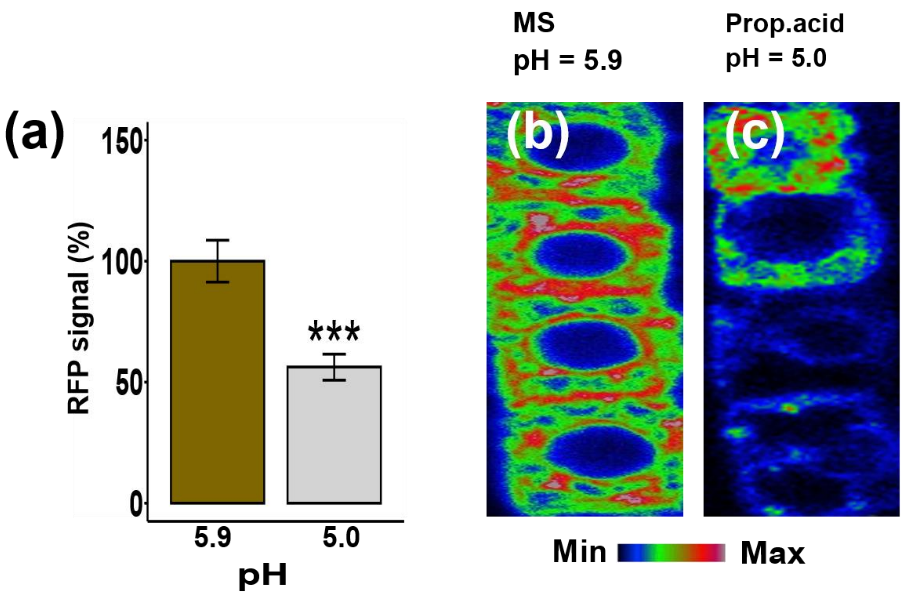


**Figure S4.** The luminal HDEL-RFP is quenched with PM permeable propionic acid.

(a) Quantified RFP signal. The RFP signal in the MS (control) treatment is plotted as 100%. The asterisks indicate significant differences in comparison to the control MS treatment (*** P < 0.001, Student's t-test). The significantly lower RFP signal after treatment with the propionic acid indicates that the luminal HDEL-RFP is quenched by the acidic pH. Error bars represent SEM from the total number of seedlings analysed from three biological experiments (n > 12 per single experiment). Scale bar = 10µm. (b-c) Images of color-coded root cells to better visualize the RFP intensity changes after the acidic treatment. Scale bar, 10 µm.

### Supplementary information

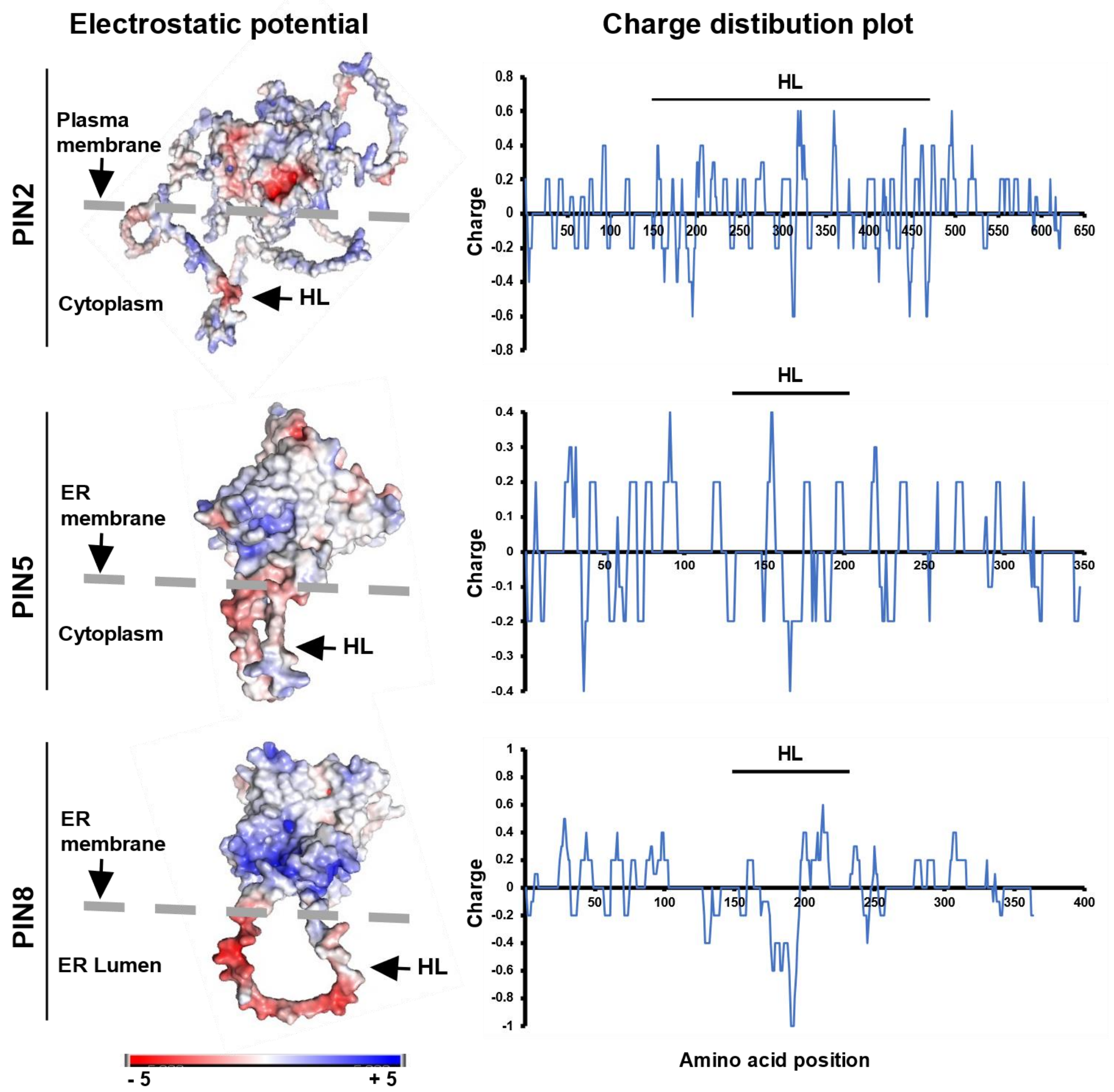


**Figure S5.** Amino acid charge distribution in PIN5 and PIN8 indicated by electrostatic potential and charge distribution plot. The three-dimensional structure of the proteins was obtained from AlphaFold database, and the surface electrostatic potential was displayed using PyMol software. Positive potential is highlighted in blue and negative potential is indicated in red. The charge distribution plot was retrieved through EMBOSS webserver. The region of the hydrophilic loop (HL) is indicated by the dash line above the x-axis. The positively charged residues are distributed in accordance with the “positive inside rule” and consequently the hydrophilic loops of PIN2 and PIN5, as verified here are positioned in the cytoplasm while the negatively charged amino acids in PIN8 HL are consistent with the luminal localisation of the HL.

### Supplementary information

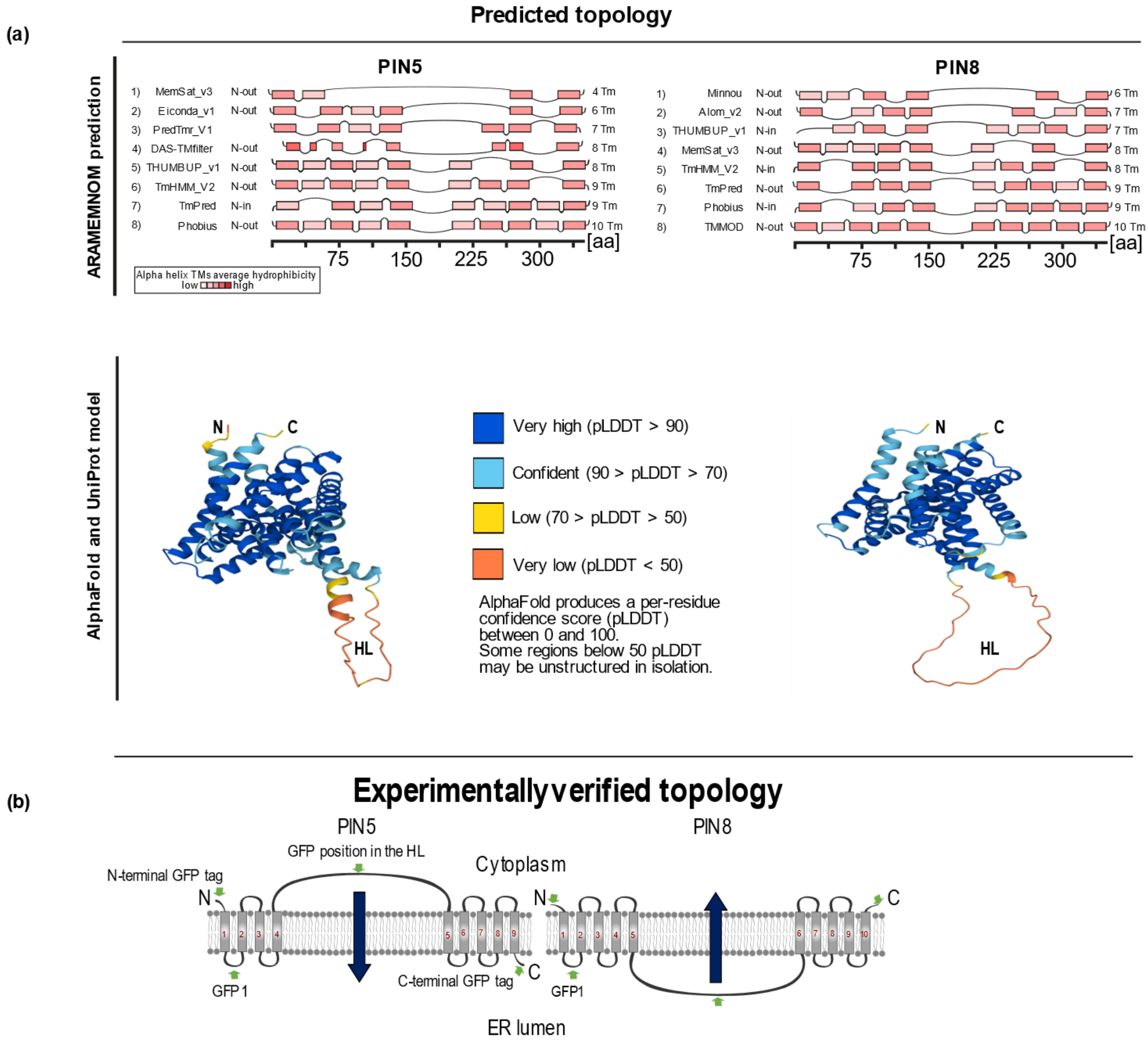


**Figure S6.** Predicted vs. experimentally verified topology of PIN5 and PIN8. (a) The PIN5 and PIN8 predicted membrane topologies obtained from the ARAMEMNOM, and the 3D structure obtained from AlphaFold database. (b) The experimentally verified membrane topology of PIN5 and PIN8. The smaller green arrows indicate the position of GFP insertion. The bigger blue arrows indicate the possible auxin transport direction of the two proteins across the ER membrane.

### Supplementary information

**Table S1.** The lists of primers used to generate the constructs

| Name of the construct or insert | | Primer name | Primer sequence |
|---|---|---|---|
| PIN2 promoter | | Pro_FP | GGGTGCAAGGATATCATTACCAGTACCG |
| | | Pro_RP | GGGTTTGATTTACTTTTTCCGGCGAGAG |
| N-terminal GFP fusion | GFP | eGFP-attB1-FP | AAAAAAGCAGGCTTCAACATGGTGAGCAAGG |
| | | eGFP-EcoRI-R | CGTCGCGAATTCCTTGTACAGCTCGTCC |
| | PIN8:GFP | P8_EcoRI_FP | CATAGCGAATTCATGATCTCCTGGCTCGATATC |
| | | P8_attB2_RP | CAAGAAAGCTGGGTCTCATAGGTCCAATAGAAAATAATATGC |
| | PIN5:GFP | P5_EcoRI_FP | GCACCGCTCGGAATTCATGATAAATTGTGGAGAT |
| | | P5_attB2_RP | CAAGAAAGCTGGGTCTCAATGAATAAACTCCAGAGCTGC |
| C-terminal GFP fusion | PIN5:GFP | attB1_P5_FP | GGGGACAAGTTTGTACAAAAAAGCAGGCTTCACCATGATAAATTGTGGAGAT |
| | | attB2_P5_RP | GGGG AC CAC TTT GTA CAA GAA AGCTGGGTCATGAATAAACTCCAGAGCTGC |
| | PIN8:GFP | attB1_P8_FP | GGGGACAAGTTTGTACAAAAAAGCAGGCTTCAACATGATCTCCTGGCTCGATATC |
| | | attB2_P8_RP | GGGG AC CAC TTT GTA CAA GAA AGCTGGGTCTAGGTCCAATAGAAAATAATATGC |
| | Fragment 1 | attB1_P5_FP | AAAAAAGCAGGCTTCACCATGATAAATTGTGGAGAT |

### Supplementary information

| | | | |
|---|---|---|---|
| PIN5 GFP 1 | | XbaI.P5_Fr 1_RP | GCACGGAGAGCGTCTAGAGAGACGGTTTATAG |
| | Fragm ent 2 | EcoRI.P5_F r2_FP | CGGGGGATTATGGAATTCGTTTGCTATTTCACCCTG |
| | | P5_attB2_R P | CAAGAAAGCTGGGTCTCAATGAATAAACTCCAGAGCT GC |
| PIN8 GFP 1 | Fragm ent 1 | attB1_P8_F P | AAAAAAGCAGGCTTCAACATGATCTCCTGGCTCGATA TC |
| | | XbaI.P8_Fr 1_RP | GCACGGAGAGCGTCTAGATGAGAAGAGCTTTAG |
| | Fragm ent 2 | EcoRI.P8_F r2_FP | GTCACCATCGAATTCCCCGAACAATGC |
| | | P8_attB2_R P | CAAGAAAGCTGGGTCTCATAGGTCCAATAGAAAATAA TATGC |
| GFP | | eGFP-XbaI-F | TCGGAGTCTAGAATGGTGAGCAAGGGCG |
| | | eGFP-EcoRI-R | CGTCGCGAATTCCTTGTACAGCTCGTCC |
